# Novel Intronic Non-Coding RNAs Contribute to Maintenance of Phenotype in *Saccharomyces Cerevisiae*

**DOI:** 10.1101/033076

**Authors:** Katarzyna B Hooks, Samina Naseeb, Sam Griffiths-Jones, Daniela Delneri

## Abstract

The *Saccharomyces cerevisiae* genome has undergone extensive intron loss during its evolutionary history. It has been suggested that the few remaining introns (in only 5% of protein-coding genes) are retained because of their impact on function under stress conditions. Here, we explore the possibility that novel non-coding RNA structures (ncRNAs) are embedded within intronic sequences and are contributing to phenotype and intron retention in yeast. We employed *de novo* RNA structure prediction tools to screen intronic sequences in *S. cerevisiae* and 36 other fungi. We identified and validated 19 new intronic RNAs via RNAseq and RT-PCR. Contrary to common belief that excised introns are rapidly degraded, we found that, in six cases, the excised introns were maintained intact in the cells. In other two cases we showed that the ncRNAs were further processed from their introns. RNAseq analysis confirmed higher expression of introns in the ribosomial protein genes containing predicted RNA structures. We deleted the novel intronic RNA structure within the *GLC7* intron and showed that this predicted ncRNA, rather than the intron itself, is responsible for the cell’s ability to respond to salt stress. We also showed a direct association between the presence of the intronic ncRNA and *GLC7* expression. Overall, these data support the notion that some introns may have been maintained in the genome because they harbour functional ncRNAs.

## INTRODUCTION

There are three main theories regarding the origin of introns: ‘Introns Late’, ‘Introns Early’ and ‘Introns First’ (Jeffares *et al*. 2006). The ‘Introns Late’ and ‘Introns Early’ theories suggest that introns arose within the eukaryotic lineage, and before the Prokaryota-Eukaryota split, respectively, whereas ‘Introns First’ implies that these noncoding sequences appeared before protein-coding genes. Introns are maintained, lost or gained with different rates within different eukaryotic lineages. The recent evolution of the yeast genome has seen widespread intron loss, with the result that only 5% of *Saccharomyces cerevisiae* genes contain introns (Dujon 2006). It has been proposed that small organisms with a large effective population size, such as *S. cerevisiae*, have seen deleterious introns gradually eliminated from the genome (Lynch 2002). This raises the question of why some introns are maintained in the yeast genome and, furthermore, if they are functionally relevant.

Recent studies indicate that *S. cerevisiae* introns increase fitness under stress (Parenteau *et al*. 2008), aid ribosome assembly, and regulate expression of the paralogous copy of the gene (Parenteau *et al*. 2011). Introns may contain regulatory sequences and structures that affect splicing or expression of their host genes. In the *HAC1* transcript, short RNA hairpins define the intron boundaries (Sidrauski and Walter 1997). Splicing of the *YRA1* mRNA regulates its export from the nucleus (Preker *et al*. 2002; Rodriguez-Navarro *et al*. 2002). Described RNA structures in pre-mRNA transcripts of *RPL30* (LI *et al*. 1996) and *RPS14B* (Fewell and Woolford 1999) regulate splicing of their host genes. Intronic hairpins in *RPL18A* and *RPS22B* pre-mRNAs are recognized by RNase III Rnt1p and promote mRNA degradation (Danin-Kreiselman *et al*. 2003). The *RPS17B* intron contains a RNA-based splicing enhancer that physically decreases the distance between the splice sites (Rogic *et al*. 2008). RNA structures in the introns of *RPS9A* and *RPS9B* transcripts are vital components of an autoregulatory circuit (Plocik and Guthrie 2012).

The cases described above are examples where introns function *in cis*; intronic sequences can also act *in trans*. In vertebrates, introns frequently harbor functional noncoding RNAs (ncRNAs). Tiling arrays and deep RNA sequencing have revealed the existence of many novel intronic transcripts, most of which have no known function (Cheng *et al*. 2005; Mercer *et al*. 2008). In addition, a recent study searching for RNAs bound by the chromatin-modifying polycomb complex raised the possibility that intronic ncRNAs can be used to guide chromatin modifications that influence gene expression in a manner analogous to some long intergenic ncRNAs (Guil *et al*. 2012). In vertebrates, there are also well-characterized examples of intronic ncRNAs such as tRNAs, small nucleolar RNAs (snoRNAs), and microRNAs (Kim and Kim 2007). However, in *S. cerevisiae*, well-known intronic ncRNAs classes are not prevalent: microRNAs are non-existent due to the loss of pre-miRNA processing enzymes (Drinnenberg *et al*. 2009), intronic snoRNAs have been mostly ‘de-intronized’ (Mitrovich *et al*. 2010) and all tRNAs are present outside introns. Even though many ncRNAs have been found in *S. cerevisiae*, such as stable unannotated transcripts (SUTs) and cryptic unstable transcripts (CUTs) (Wu *et al*. 2012), only 2% of the instances of these classes overlap with introns (Xu *et al*. 2009). It is commonly believed that splicing in higher eukaryotes increases protein diversity by providing multiple mRNAs from a single locus. However, alternative splicing in *S. cerevisiae* has been shown only for transcripts from the genes *SRC1* (Davis *et al*. 2000; Grund *et al*. 2008), *PTC7* (Juneau *et al*. 2009) and *MTR2* (Davis *et al*. 2000; Preker *et al*. 2002). Since alternative splicing in *S. cerevisiae* is rare and there has been a massive reduction in the number of typical intronic ncRNAs, we hypothesize that functional yeast introns may have been retained because they contain novel ncRNA genes or pre-mRNA RNA structures.

In order to discover potential functional RNA structures within introns in *S. cerevisiae*, we performed a computational screen for novel structures using intron orthologs from 36 fungal species and employing three *de novo* RNA structure prediction tools. The screen identified 19 introns containing potential RNA structures, and we validated the expression and processing of a subset by RT-PCR. We showed that six introns tested are maintained in the cell after splicing and two contain novel ncRNAs. A novel ncRNA within the *GLC7* intron, rather than the whole intron (Juneau *et al*. 2006; Parenteau *et al*. 2008), is responsible for cell’s ability to respond to salt stress, by altering the gene expression.

## RESULTS

**Predictions of RNA structure within introns**: Since a predicted secondary structure of a single sequence is not generally sufficient to distinguish between a functional RNA and random sequences (Rivas and Eddy 2000), most RNA prediction methods require multiple homologous sequences. Thus our first step in RNA prediction was to identify orthologs of *S. cerevisiae* introns in other fungi. We searched for orthologs of intron-containing host genes in fully sequenced genomes and then for corresponding introns in those genes. In 36 fungal genomes, we were able to identify at least 2 orthologs for 281 introns and at least one ortholog for 305 introns (see Fig 1A). Only the intron of YDR535C does not have an ortholog in any of the species searched, but annotation of this sequence as a gene is dubious. The vast majority of introns are conserved only in the *Saccharomyces* ‘sensu stricto’, and we identify fewer than 4 orthologs for each intron on average. In contrast, orthologs of yeast introns containing known intronic snoRNAs are found in a wider range of fungal genomes, having on average 9.6 orthologs (see Fig 1A).

**Figure 1.**
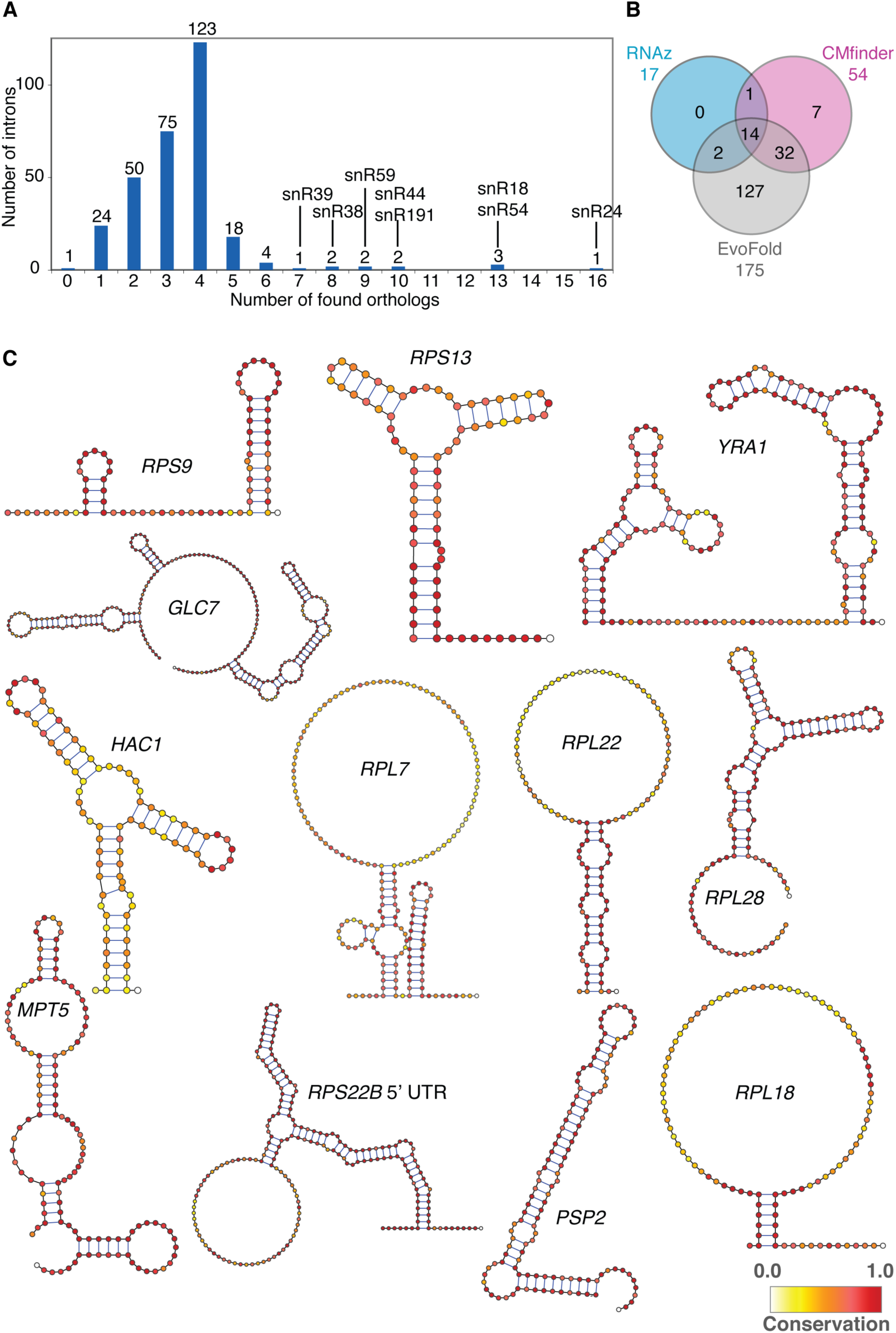
– Predicted RNA structures. - A Distribution of the number of orthologs identified for *S. cerevisiae* introns among 36 species searched. Introns containing known snoRNAs are indicated by name.
- B Venn diagram showing the number of introns with predicted RNA structures by different programs. RNAz, CMfinder and EvoFold predicted structures in 14 common introns.
- C Predicted consensus RNA structures. The name of the corresponding gene is shown next to each structure. In the case of duplicated ribosomal gene introns where both paralogous introns share a similar structure, the standard name of the gene is given. Structure images were prepared using VARNA (Darty *et al*. 2009).

Orthologous intronic sequences were used to predict novel RNA structures using three independent computational methods: CMfinder, RNAz and EvoFold. Each of these approaches yielded a very different number of predicted RNA structures in introns. RNAz, CMfinder and EvoFold identified putative conserved RNA structures in 17, 54 and 175 introns, respectively. We found 14 structures in the intersection of all three approaches, within the only intron of each of *GLC7*, *HAC1*, *IMD4*, *MPT5*, *RPL18A*, *RPL18B*, *RPL22B*, *RPL28*, *RPS9A*, *RPS9B* and *RPS13*, in the first intron of *RPL7A* and in both introns of *RPS22B* (Fig 1B and Supporting Information 1).

For further bioinformatical and experimental analysis, we decided to focus on the introns that had structures predicted by all three programs, together with their paralogs (*i.e*. introns in *RPL22A*, *RPL7B*), and 3 introns with high prediction scores in at least two out of three approaches (*i.e*. introns in *NOG2*, *YRA1* and *PSP2*). We used the INFERNAL software (Nawrocki *et al*. 2009) and the Rfam library of covariance models to search the intron sequences for known ncRNA classes (Gardner *et al*. 2010). Besides the known snoRNAs, our other high-scoring RNA predictions do not resemble any previously known RNA families. We also used an iterative procedure combining covariance model searches using the INFERNAL package and manual inspection of multiple sequence alignments to identify additional homologs of our predicted structures in more distant species (Table I and Fig 1C). We find that the known snoRNAs are very well conserved among Fungi: snR191 orthologs were found in all Saccharomycotina, snR44 in Saccharomycotina and Pezizomycotina, while snR54 is present in some Metazoan genomes as well as Fungi. The previously known *HAC1* intron-exon structure is also conserved in Fungi and Metazoa (Hooks and Griffiths-Jones 2011). Iterative INFERNAL searches for homologs of the predicted structures in the introns of *RPL18*, *RPL22*, *RPL7* and *RPS9* allowed us to extend the conservation to the *Saccharomyces* and *Candida* clades. The identification of highly similar short motifs in the alignments of *RPL7*, *RPL18*, *RPL22*, *RPL28* and *RPS9* introns demonstrated that predicted structures within introns are well-conserved in *Saccharomyces* ’sensu stricto’. The high sequence conservation means that relatively few compensatory mutations support these conserved predicted structures. Only in the case of the intron of *RPS13* is there evidence for maintenance of the RNA structure through multiple compensatory mutations.

**Table I.** Conservation of RNA structure predictions.

| # | Locus | Gene | Conservation |
| --- | --- | --- | --- |
| 1 | YBR189W<br>YPL081W | *RPS9B*<br>*RPS9A* | Saccharomycetaceae and *Candida* sp. |
| 2 | YDR064W | *RPS13* | Saccharomycetaceae |
| 3 | YDR381W | *YRA1* | Saccharomycetaceae |
| 4 | YER133W | *GLC7* | *Saccharomyces* 'sensu stricto' |
| 5 | YFL031W | *HAC1* | Fungi and Metazoa |
| 6 | YFL034C-A<br>YLR061W | *RPL22B*<br>*RPL22A* | Saccharomycetaceae |
| 7 | YGL076C<br>YPL198W | *RPL7A*<br>*RPL7B* | Saccharomycetaceae and *Candida* sp. |
| 8 | YGL103W | *RPL28* | *Saccharomyces* 'sensu stricto' |
| 9 | YGL178W | *MPT5* | *Saccharomyces* 'sensu stricto' |
| 10 | YLR367W<br>5' UTR | *RPS22B* | Saccharomycetaceae except *L. lactis* |
| 11 | YLR367W<br>snR44 | *RPS22B* | Saccharomycotina and Pezizomycotina |
| 12 | YML017W | *PSP2* | *Saccharomyces* 'sensu stricto' |
| 13 | YML056C<br>snR54 | *IMD4* | Saccharomycetales and Diptera |
| 14 | YNL301C<br>YOL120C | *RPL18B*<br>*RPL18A* | Saccharomycetaceae and *Candida* sp. |
| 15 | YNR053C<br>snR191 | *NOG2* | Saccharomycetaceae and *Candida* sp. |

**Experimental detection of the predicted intronic ncRNA**: In order to determine if the predicted intronic RNAs are expressed and maintained in the cell, we performed RT-PCR on total and low molecular weight cDNA from haploid BY4741 wild type *S. cerevisiae* strain. We designed three sets of specific primers to amplify the fragment of intron with the predicted structure or snoRNA, the entire intron, and part of the exons flanking the intron. As a positive control for the PCR, we used the genomic DNA to amplify products with the primer pairs described above in the snR44, snR191 and snR54 snoRNA genes. Negative controls for all PCR experiments were performed with no template added to the reactions.

If a *bona fide* ncRNA product is processed from an intron, we expected to obtain bands corresponding to the spliced mRNA of the host gene, and also for the region of the predicted structure, but not the larger product from the complete intron. This pattern was observed for the *RPS22B* intron harboring the known snoRNA snR44 and for two other introns with predicted structures, namely the intron of *GLC7* and the first intron of *RPL7B*, thus confirming these sequences as novel ncRNAs (Fig 2A, Fig S1). A similar pattern was also observed for *MPT5* although its expression was very low (Fig 2A, Fig S1C). The RT-PCR corresponding to snR191 in *NOG2* displayed a pattern indicative of complete splicing of the host mRNA, but also showed the maintenance of the complete intron and of the predicted ncRNA. We refer to these sequences as ‘introns maintained’ after splicing as opposed to ‘introns retained’ in the pre-mRNA. We observed a similar pattern of correct splicing with intact intron and ncRNA maintenance for our predicted structures in the introns of *RPS13, RPS9B, RPL7A*, *RPL22A* and in the 5’ UTR intron of *RPS22B* (Fig 2B and Fig S2-S3). These data show that a mixed population of mature ncRNAs and spliced but unprocessed introns are present in the cell. We also found that the mRNA transcripts of *IMD4* (containing snR54 in its intron), *PSP2, RPS9A, RPL28, RPL22B, RPL18A, RPL18B* and *YRA1* were present in both spliced and unspliced forms, supporting the existing evidence that intron retention is the most common case of alternative splicing in yeast (Plass *et al*. 2012) (Fig 2C, Fig S4-S5). We found no evidence of splicing of *HAC1* under the specific experimental conditions we tested (see Fig S6A). As negative controls, we conducted the same RT-PCR analysis on six intron-containing genes that had no predicted RNA structures. None of the six displayed patterns consistent with ncRNAs processed from the introns. The genes YBR219C and *BMH2* did not appear to be expressed and the four ribosomal protein genes *RPL27A*, *RPS27B*, *RPS16A* and *RPS19B* showed the intron-maintained pattern (Fig S6B-G). RNAseq data show higher expression of introns in ribosomal protein genes that contain predicted RNA structures.

**Figure 2.**
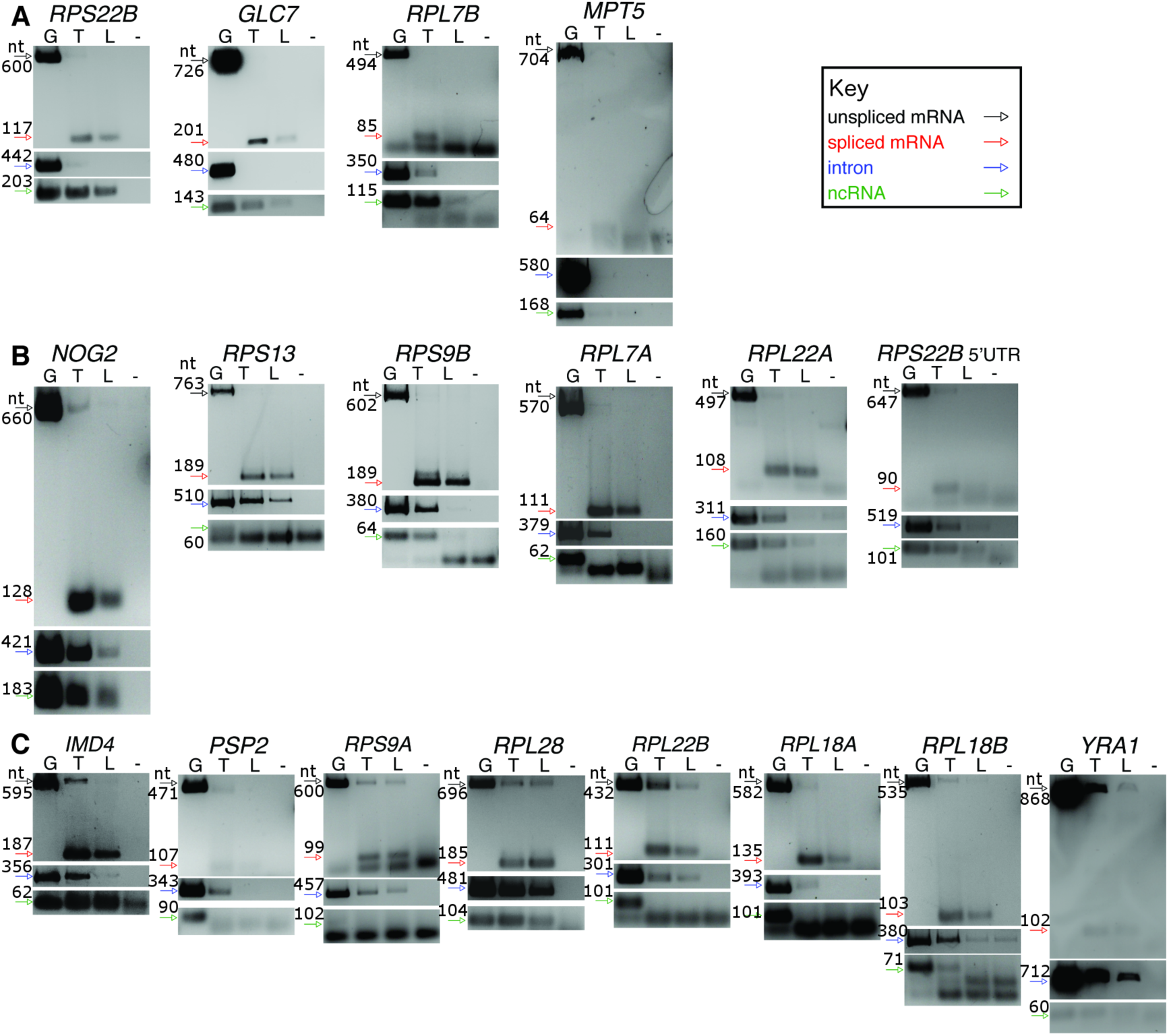
– RT-PCR confirmation of intron fates. - A Agarose gel confirming ncRNAs expressed from introns and maintained in the cell.
- B Agarose gel confirming ncRNAs expression accompanied by complete mRNA splicing.
- C Agarose gel confirming ncRNAs expression accompanied by alternative splicing. Arrows indicate the expected size of the PCR products according to the key. Lane designations for DNA templates: G, Genomic DNA (positive control); T, total cDNA; L, low molecular weight enriched cDNA; (-), no template negative control. With the exception of *GLC7* ncRNA, which was run on a separate gel, the images of mRNAs, introns and ncRNAs for each gene were cropped from the same agarose gel picture with brightness and contrast applied equally across the entire image (full images available in Figures S1-S6). For small size PCR products, cropping included primer dimers.

We also validated the presence of expressed intronic sequences in the cell by deep sequencing of total RNA extracted from *S. cerevisiae* BY4741. We counted all reads overlapping introns and normalized by the intron length and the total number of reads mapped to obtain values of **r**eads **p**er **k**B per **m**illion mapped (RPKM). Median intron expression was 15.3 RPKM. We observed that 17 of 19 introns with predicted structures have evidence of expression (more than 150 reads or 20 RPKM), seven of which fell within the 90th percentile of intron expression.

Since a third of all introns are found in highly expressed ribosomal protein (RP) genes, we next measured the relative expression levels of maintained introns in RP that contain or do not contain predicted structures. Since the mRNA levels of the genes hosting predicted intronic structures appeared to be greater than the average mRNA amount for the entire set of intron-containing genes, we also normalized the intron expression levels by the level of their host gene transcript. Interestingly, expression of RP introns with predictions was significantly higher compared with all RP introns before and after normalizing for host gene expression (median RPKM 65.5 compared to 33.0, Mann–Whitney U test p-value: 0.006 and median normalized expression 0.11 compared to 0.03, Mann–Whitney U test p-value: 0.014, Fig 3A).

**Figure 3.**
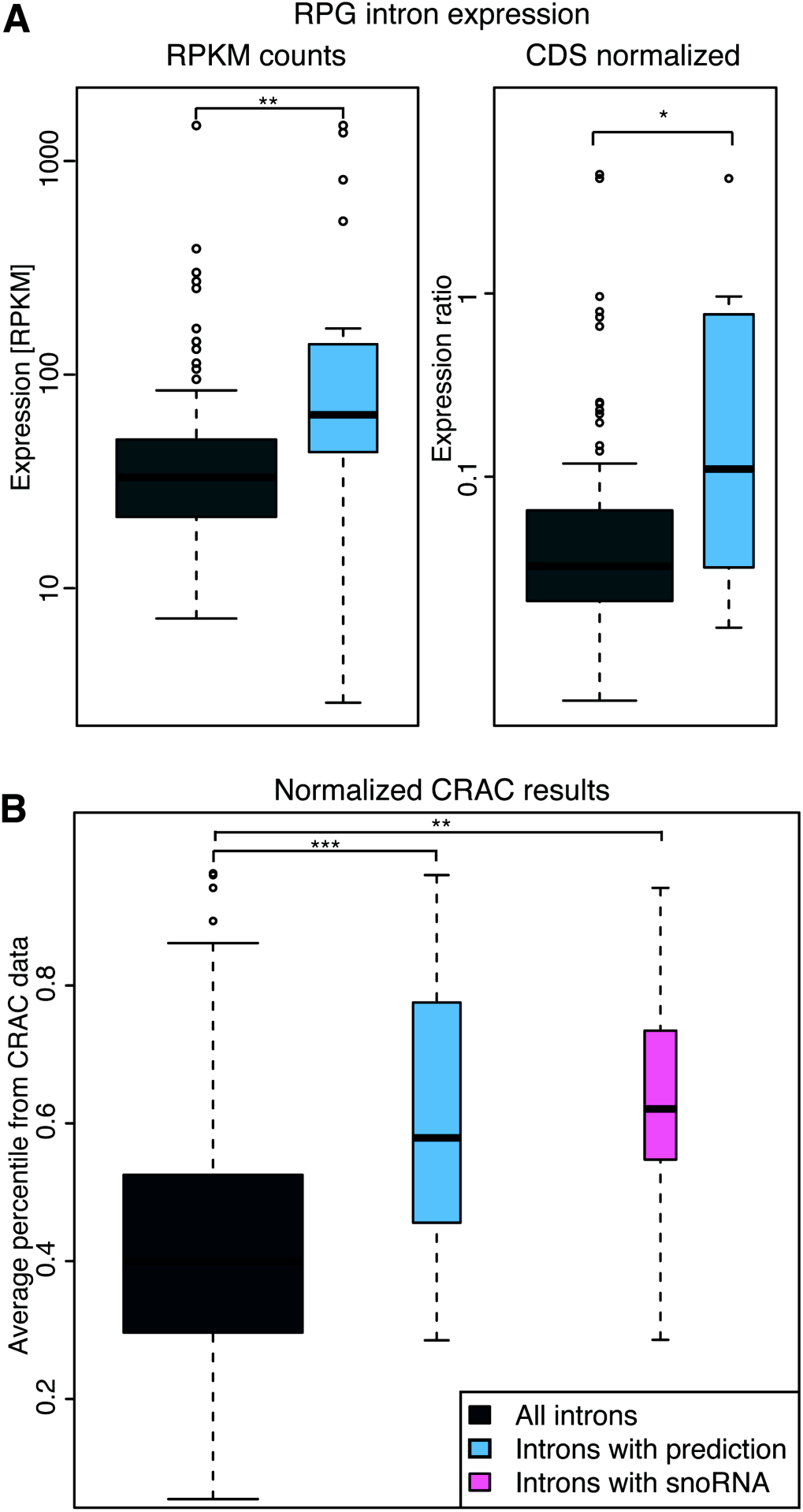
Properties of loci containing intronic novel RNA predictions or snoRNAs. - A Box plots showing ribosomal protein intron transcript levels without normalization in RPKM and with normalization to host gene transcript levels. Values are shown for all RPG introns and twelve RPG introns with predictions
- B Normalized average percentile of reads bound to protein targets derived from sixteen CRAC experiments taken from Schneider *et al*. (2013). Values are shown for all introns, the nineteen introns with predictions and the eight introns containing snoRNAs. The levels of significance for the Mann-Whitney U tests are represented as follows: ***: p < 10^−3^; **: p <0.01; *: p <0.05.

**Introns with predicted RNA structures are more likely to be targeted by the exosome**: We hypothesize that introns with RNA structures either contain novel noncoding RNAs or are involved in the regulation of pre-mRNA splicing; in both cases, the host transcript would be expected to associate with the exosome complex. We analyzed the data of Schneider *et al*. (Schneider *et al*. 2012), who used *in vivo* RNA crosslinking (CRAC) of exosome components to show that noncoding RNAs, snoRNAs, pre-tRNAs and pre-mRNAs are the most prominent exosome targets. The Schneider *et al*. dataset (GEO accession GSE40046) contains deep sequencing reads from sixteen separate *in vivo* crosslinking experiments to the tagged protein components of the exosome. When we normalized the number of reads of each intron-containing gene from GSE40046 by the average number of reads for this intron-containing gene from our sequencing, we observed that the set of genes with both known snoRNA host genes and our novel RNA predictions appears to be targeted by the exosome machinery more than expected (our predictions, Mann–Whitney U test, p = 2.44×10^−4^; snoRNAs, p = 0.005; Fig 3B). This suggests that the introns of interest are either retained in pre-mRNAs or contain noncoding RNAs. Taken in conjunction with our RT-PCR data, the CRAC data indicated that the transcripts of *PSP2*, *RPS9A*, *RPL18A* and *RPL18B* retain introns, and as a result their pre-mRNAs are targeted for exosome-mediated degradation. In contrast, the presence of novel ncRNA was indicated for the *MTP5* intron.

**Function of the *GCL7* intronic ncRNA**: The intronic sequence in *GLC7* was recently shown to play a role in the cellular response to osmotic pressure (Parenteau *et al*. 2008). We therefore investigated further the putative ncRNA derived from this intron. Firstly, we used Northern hybridization to confirm the RT-PCR analysis of expression of the ncRNA; and secondly we employed primer walking to define the 5‘ and 3‘ ends of the ncRNA. We were able to specifically define *GLC7* ncRNA boundaries, which are different to those of the previously annotated CUT568 residing on the opposite strand (Fig 4).

**Figure 4.**
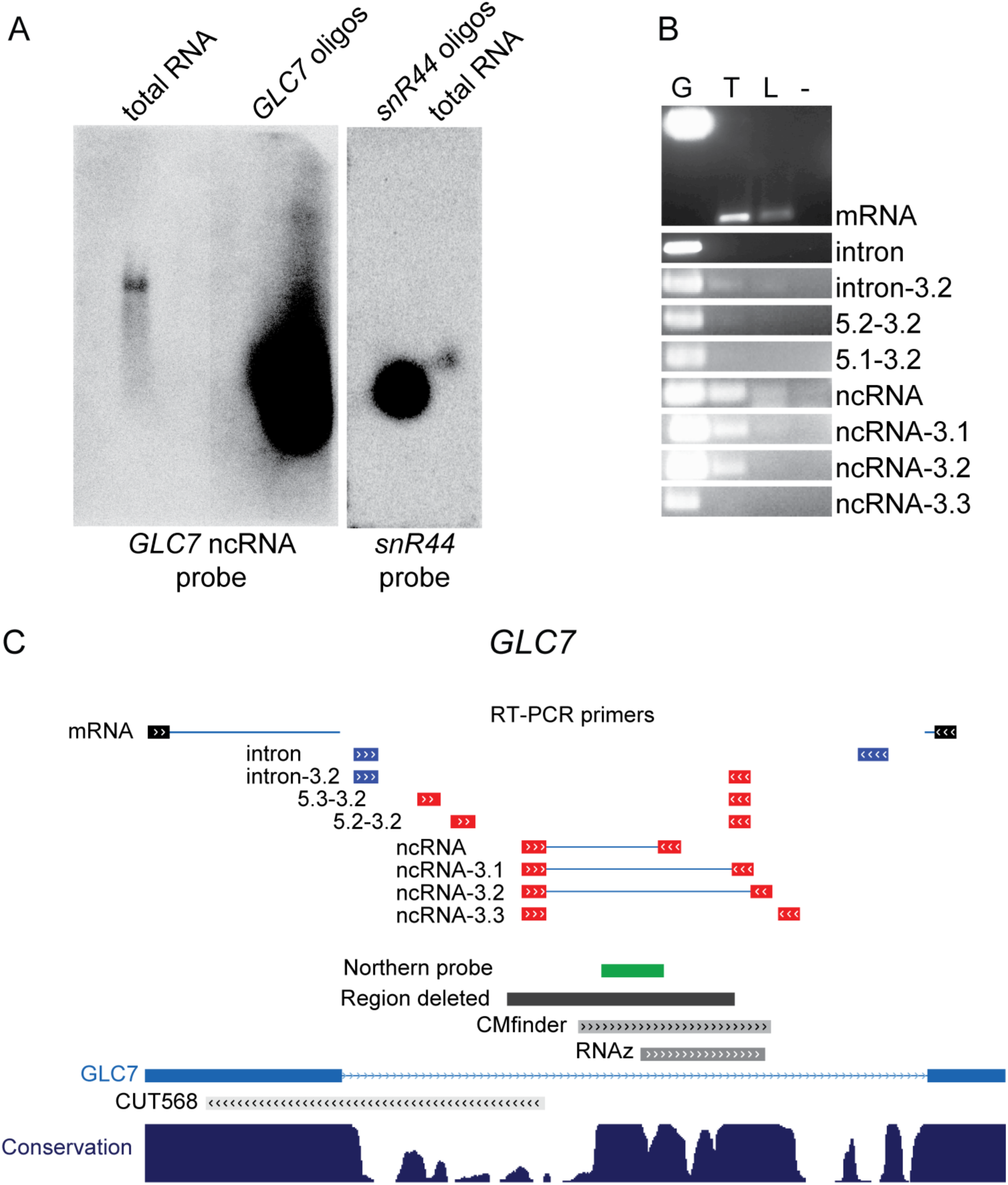
– Characterization of the size and expression of the ncRNA within *GLC7* intron. - A Northern blot of *S. cerevisiae* size fractionated RNA and RNA oligos mimicking *GLC7* intron or *snR44* probed with strand specific probes complementary to ncRNA in *GLC7* (left panel) and *snR44* (right panel).
- B RT-PCR gels for *GLC7* with primers listed. Gel annotation: G, genomic DNA; T, total cDNA; L, low molecular weight enriched cDNA; -, no template negative control.
- C Data uploaded into the UCSC genome browser for sequence annotation and data visualization presenting annotated *GLC7* intron. Primer names used for RT-PCR are listed next to black, blue and red boxes indicating their position in respect to the gene annotation below. Lines joining primers mark the amplified sequences, the regions targeted for deletion by the loxP method are symbolized by black box, location of the strand-specific Northern probe in green and regions with the putative structure predicted by RNAz and CMfinder are shown in grey. All features map directly onto the fragment of the gene structure diagrams. The bottom panel represents the degree of conservation of gene regions among seven yeast species.

To study the phenotype of the ncRNA we used two approaches: deletion and alteration of the ncRNA structure. We deleted two regions in the intron of *GLC7* via PCR-mediated gene deletion and the cre-loxP system: an intronic region overlapping with CUT568 but not with the ncRNA (negative control deletion mutant) and the intronic region corresponding to the ncRNA (*GLC7* ncRNA deletion mutant). For the alteration, we inserted 139 bp in the middle of the predicted ncRNA to modify its structure (*GLC7* ncRNA insertion mutant, see Methods). Previously it was shown that replacing the *GLC7* gene with its cDNA decreases the cell viability in NaCl stress (Juneau *et al*. 2006; Parenteau *et al*. 2008). In order to determine if this defect was due to the ncRNA structure rather than the intron in itself or the CUT568, we tested all engineered strains with the mutated *GLC7* intron in F1 medium and in F1 medium containing 0.9 M NaCl (Fig 5A). In F1 supplemented with 0.9 M NaCl, we observed a significant difference in growth, as estimated by the area under the growth curve (Norris *et al*. 2013), for both *GLC7* ncRNA deletion and insertion mutants (Student’s t-test, p = 1.4 × 10^−9^ and p = 1.88 × 10^−7^, respectively) compared with the wild type or the negative control deletion (p = 2.44 × 10^−9^ and p = 9.67 × 10^−9^ respectively, Fig 5B). Specifically, we found that in the F1 + 0.9 M NaCl media the lag phase (λ) is longer in the *GLC7* ncRNA deletion and insertion mutants. Moreover, their maximum growth rate (µ) and the final biomass after 48 hours (A) are decreased, whereas for the negative control deletion mutant those parameters are the same as the wild type (Fig S7).

**Figure 5.**
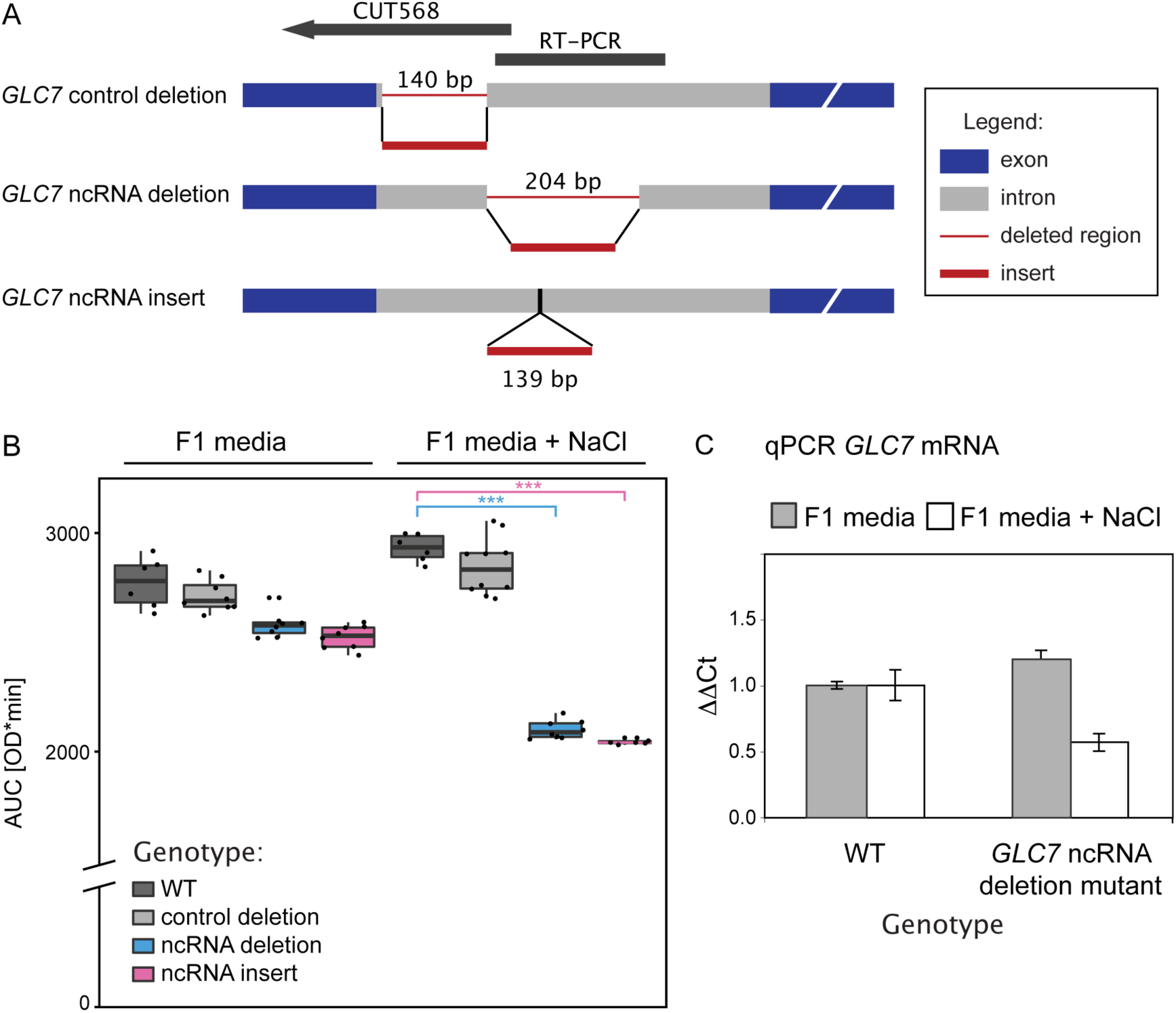
– Effects of *GLC7* intron mutation. - A Schematic representation of intron mutants used for phenotype studies.
- B The *GLC7* ncRNA deletion and *GLC7* ncRNA insertion mutants disrupting the structure sequence of *GLC7* intron were compared with the wild type strain and the intronic negative control in F1 medium containing 0.9 M NaCl. Values in the box plots present the means of the area under the growth curve (AUC) as determined by R pracma package. Significant p-values obtained from Student’s t-tests are indicated (***: p < 10^−6^).
- C Expression levels (average expression with standard error bars) of *GLC7* mRNA in the WT and in the *GLC7* ncRNA deletion mutant grown in F1 and F1 + 0.9M NaCl media, assessed by real time PCR.

We further verified the phenotypic difference of the mutant using one-to-one competition experiments. The wild type BY4742 strain and the *GLC7* deletion mutant were grown together in F1 medium supplemented with 0.9 M NaCl and the composition of the population was analyzed after 37 generations. A significant drop in relative amount of the mutant strains was detected with just over 25 % of total cells being *GLC7* deletion mutant, instead of the expected 50 % (Student’s t-test, p = 7.30x10^−5^). Taken together, these data further indicate that the deletion of the putative ncRNA in the *GLC7* intron is responsible for the observed impairment of cell growth rate and competitive fitness during salt stress, whereas deletion of the upstream intronic region overlapping the CUT568 expressed from the opposite strand has no impact on the phenotype in salt stress.

We then looked at the expression of *GLC7* via real time PCR in both wild-type and *GLC7* ncRNA deletion mutant. The data shows that the deletion of the ncRNA in *GLC7* intron does not impair splicing and that the *GLC7* mRNA level appears to be slightly elevated in F1 media compared to that of wild type (1.21 ± 0.07). However, in salt stress the *GLC7* mRNA level of the *GLC7* ncRNA deletion mutant is reduced to 0.60 ± 0.12 of that of the wild type (Fig 5C). We suggest that the phenotypic effect of the ncRNA knockout under salt stress may be due to the decreased expression of *GLC7* in the *GLC7* ncRNA deletion mutant.

## DISCUSSION

Using multiple computational methods, we predicted stable and conserved RNA structures in 19 introns, 12 of which are present in ribosomal protein genes. By RTPCR, we validated the presence of the predicted ncRNAs and, in several cases, of the whole introns. Our predicted RNA structures include in the 5’ UTR intron of *RPS22B* (Fig 2B) and in the intron of *RPL18A* (Fig 2C). Both introns have been previously reported to trigger RNase III-mediated mRNA degradation by Rnt1p (DaninKreiselman *et al*. 2003). We also predicted and validated the expression of putative ncRNAs in the introns of *RPS9A* and *RPS9B* (see Fig 2B), and these introns were previously shown to regulate expression of both their host genes and their paralogs (Plocik and Guthrie 2012). In another study on the effects of intron deletion (Parenteau *et al*. 2011), 11 RP introns besides *RPS13* were shown to regulate the expression of the host gene or its paralogous copy, change cell sensitivity to drugs or alter the competitive fitness when deleted. Our RNAseq data show that introns in RP genes containing RNA structures are significantly more expressed than the introns in RP genes lacking a predicted ncRNA. Among the seven non-RP proteins with high-scoring predictions, only the *PSP2* introns have no previously suggested function. *HAC1* and *YRA1* introns contain structures that regulate their own splicing and *IMD4* and *NOG2* contain snoRNAs that guide chemical modification of other RNAs, whereas *GLC7* and *MPT5* introns appear to be required for stress tolerance (Parenteau *et al*. 2008). Our data raise the possibility that, at least in some cases, a processed ncRNA is responsible for the biological function, rather than the intron itself.

Our experimental characterization and follow-up analysis has allowed us to generate hypotheses regarding the mechanism of function of specific sequences. For example, the *GLC7* intron was previously shown to mediate the response to salt stress (Parenteau *et al*. 2008) and we now demonstrate that the factor responsible for the biological function is an intronic ncRNA sequence with a discreet RNA structure (see Fig 5). Furthermore, characterization of the stable ncRNA by RT-PCR suggests that the predicted structure functions *in trans* after removal from an intron lariat to enhance the expression of the *GLC7* gene. Similarly, a recent study in human cancer cells showed that the *SMYD3* intron may negatively regulate the expression of the *SMYD3* gene *in trans* by interactions with a *EZH2* protein, a core component of Polycomb Repressive Complex 2 (Guil *et al*. 2012). New techniques, such as *in vivo* RNA crosslinking to protein complexes coupled with next-generation sequencing, can aid the discovery of novel pathways involving all types of ncRNA, including those encoded in introns. Our analysis of exosome target data presented by Schneider *et al*. (2013) indicates genes with intronic RNA structure predictions are more likely to be transcriptionally regulated or contain novel ncRNAs.

With the exception of a few snoRNAs, *S. cerevisiae* introns do not appear to contain classical intronic ncRNAs. Although the function of most intronic RNAs in higher eukaryotes is still unknown, the evidence of tissue-specific expression (Louro *et al*. 2008) and binding to protein complexes known to promote epigenetic modifications (Guil *et al*. 2012) indicates that intronic transcripts may have specific functions, rather than arising from spurious transcription or slow pre-mRNA turn-over. Our results provide clear evidence for the function of the *GLC7* intronic ncRNA in an intron-poor single-celled fungal species.

Our work, and that of others mentioned here, raises interesting questions about the general nature of intron stability post-splicing. Introns removed by splicing have been thought to be rapidly degraded and the main focus of intron biology has concerned elucidation of splice sites and the arrangement of the splicing machinery. For example, it was observed that the deletion of the debranching enzyme promotes the accumulation of lariat introns (Chapman and Boeke 1991), so it was assumed that all introns are rapidly debranched and targeted for degradation in normal cells. Tiling arrays and next generation sequencing have readily shown that some intronic sequences are abundant in the cell, but they are usually dismissed as remnants of normal splicing or part of immature pre-mRNAs (Louro *et al*. 2009). Recently, the fate of introns themselves has been systematically assessed in *Xenopus tropicalis* embryos showing that 90% of intron are maintained in the nucleus post splicing (Gardner *et al*. 2012) and around 5% of genes generate lariats that are stable in the cytoplasm (Talhouarne and Gall 2014). Linear intron derived ncRNA with some similarities to snoRNAs have been found in HeLa cells and human embryonic stem cells (Yin *et al*. 2012). Most importantly, circular intronic RNAs increase the expression of their host genes as exemplified by gene *ANKRD52* (Zhang *et al*. 2013). We and others observe that many of the introns examined are retained in the premRNA or maintained in the cell after splicing. This is consistent with the observations of Coleclough and Wood (1984) who were the first to described discrete intron products processed from the mouse immunoglobin pre-mRNA. Immunoglobulin mRNAs are expressed at high levels, like the ribosomal genes tested in our study. We therefore speculate that post-splicing intron stability might be a prevalent phenomenon for highly expressed genes and that intron products may be regulators of these highly expressed gene products.

## CONCLUSIONS

We undertook a systematic approach using the well-studied *Saccharomyces cerevisiae* genome in order to identify introns with undiscovered function. Comparing intron sequences of related yeast species, we found at least 19 introns contain putative conserved RNA structures. By RNAseq and RT-PCR, we show that several of the intronic sequences containing secondary structures are not degraded after removal from pre-mRNAs. Furthermore, we show that ncRNAs embedded in introns can be directly responsible for regulating gene expression and maintaining phenotype in the intron-poor yeast. For example, a small portion of the *GLC7* intronic sequence, representing a novel ncRNA, plays an important role in the cellular response to salt stress. More generally, the cellular abundance of intron sequences from ribosomal protein genes with predicted intronic ncRNAs are significantly more highly expressed than those lacking such predictions. Overall, our data support the possibility that the presence of functional RNA structures in introns has contributed to selective intron retention in the *Saccharomycetes*.

## MATERIALS AND METHODS

**Intron alignments**: Sequences of intron-containing genes were extracted from the Saccharomyces Genome Database (SGD, http://www.yeastgenome.org/). Genes orthologous to intron-containing genes from *S. cerevisiae* were identified in 36 fungal genome sequences (Table S1) using TBLASTX with the coding gene sequence as the query, and with following settings: –E e-6 -qframe 1 -hspsepsmax 1000 -topcomboN 1. We collected the sequences of putative orthologs with at least 65% query coverage, to which 1000 bp and 300 bp of flanking sequence were added to the 5‘ and 3‘ ends of each hit. The putative ortholog gene sequences were then searched for the presence of the orthologous intron using BLASTN with options: -E 0.1 -W 3 -hspsepsmax 1000. The best hit (with the lowest e-value and confirmed by manual inspection) was retained for each intron in each species.

**RNA structure predictions**: Three structure prediction programs were used: CMfinder (v 0.2) RNAz (v 2.0) and Evofold (v 7b). Sequences of orthologous introns were used for predictions with CMfinder as described in Torarinsson *et al*. (2008). CMfinder was run with settings ‘-n 5 –m 30 -M 100’ and ‘-s 2 -n 5 -m 40 -M 100’ and identified motifs were extended using the CombMotif.pl procedure. Motifs with a composite score of r > 5 and folding free energy of < -5 kcal.mol^-1^ were considered as putative positives. For RNAz and EvoFold predictions, intron sequences were first aligned with mLAGAN. Structure prediction by RNAz was performed according to the manual (http://www.tbi.univie.ac.at/~wash/RNAz/manual.pdf). The, rnazWindow.pl script was used to slice the alignments with following options: --maxgap=0.25 --min-id=30 --max-seqs=6. RNAz was then run on the forward strand of the gapped alignments (options: --forward –g –p 0) and sequences with probability P > 0.5 were considered as putative positives. For EvoFold predictions, the required phylogenetic tree containing the species present in the intron alignments was derived by pruning the tree presented by Medina *et al*. (2011), and the subsequent structure predictions were predicted using default parameters. We employed a threshold value of 10 for the log-odds ratio of the likelihood of the region under the structure model and background model. The complete list of all predictions can be found in Supporting Information 1. To extend the phylogenetic range of the RNA predictions, BLAST and INFERNAL 1.0.2 (Nawrocki *et al*. 2009) were used to re-search all fungal genomes in an iterative process, based on the Rfam approach (Gardner *et al*. 2010) and as described previously (2011).

**Strains and media**: Intron sequence replacement strains were engineered from the BY4743 (*MAT*a/α, *his3*Δ1/*his3*Δ*1*, *leu2*Δ*0*/*leu2*Δ*0*, *lys2*Δ*0*/ *LYS2*, *MET15*/*met15*Δ*0*, *ura3*Δ*0*/*ura3*Δ*0*; 4741/4742), BY4742 (*MAT*α, *his3*Δ*1*, *leu2*Δ*0*, *lys2*Δ*0*, *ura3*Δ*0*) and BY4741 (*MAT*a, *his3*Δ*1*, *leu2*Δ*0*, *met15*Δ*0*, *ura3*Δ*0*) parental strains and maintained in YPD medium containing 2% (w/v) yeast extract, 1% (w/v) peptone and 2% (w/v) glucose. The transformants were plated on solid YPD medium with 300 mg/ml geneticin (GibcoBRL, UK) for kanMX selection and on 10 mg/ml phleomycin (Invitrogen) for pCre-ble selection. Mineral salt F1 media was prepared as described previously (Baganz *et al*. 1998) and the synthetic minimal SD medium (0.67% Bacto yeast nitrogen base without amino acids, 2% glucose) was supplemented with required amino acids appropriate for the parental strain.

**RNA extraction**: The *S. cerevisiae* strain BY4741 was grown in 500 ml rich media (YPD) in 30 ºC with shaking at 200 rpm to an absorbance of 0.5 at 600 nm. The RNA was extracted using Trizol (Invitrogen, UK), precipitated in lithium chloride (Ambion, UK), washed twice with 70% ethanol and the pellet re-suspended in dH2O. RNA concentration and quality was evaluated by measuring absorbance at 260 nm on a NanoDrop spectrometer ND-1000 (Thermo Scientific). The low molecular weight enriched RNA sample was obtained from total RNA as described in Catalanotto *et al*. (2002). Total RNA was used for RNAseq and both total and low molecular weight enriched RNA was used for RT-PCR.

**RT-PCR**: cDNA was synthesized from 2 µg RNA of either total or low-molecular weight RNA using QuantiTect Reverse Transcription Kit (QIAGEN) according to the manufacturer’s protocol. Fragments of cDNA corresponding to the predicted intronic RNA structure ('ncRNA‘), whole intron of interest ('intron‘) and exons surrounding the intron ('mRNA‘) were amplified by PCR with BIOTAQ DNA Polymerase (Bioline) according to the supplier‘s guidelines. The list of all primer sequences used can be found in Supporting Information 2. The reaction mix was comprised of 4 pmol of each primer and 20 ng of total or low molecular weight cDNA for each 10 µl of total reaction mixture. The cycling conditions were an initial denaturation for 5 min at 95°C, 35 cycles of denaturation (45 sec, 94°C), annealing (45 sec, 56°C) and elongation (90 sec, 72°C) followed with final elongation for 5 min at 72°C. Amplification of both cDNA using snR44 primers was used as a positive control. Genomic DNA extracted from BY4741 with Wizard Genomic DNA Purification Kit, PROMEGA, according to the manufacturer’s protocol was used as a positive control and water was used as a negative control. The PCR products were visualized by ethidium bromide staining on 1.2-3.5% agarose gels. For each predicted RNA structure, at least two independent PCR reactions with genomic DNA, total cDNA, low molecular weight cDNA and snR44 positive controls primers were performed in order to confirm expression.

**Northern hybridization**: 20 µg of the size fractionated RNA and 10 pmol of the oligonucleotides mimicking the intronic region of *GLC7* and snR44 complementary to the probes subsequently used for hybridization, were loaded in RNA loading dye (Fermentas) onto separate lanes of a denaturating gel containing 36.5mM MOPS, 9.1mM sodium acetate, 0.9mM EDTA, 2M formaldehyde, 0.5µg/ml ethidium bromide and 1% agarose. RNA transfer, UV-crosslinking and Northern blotting was performed as described before (Naseeb and Delneri 2012). As probes [^32^P]-ATP end-labeled antisense oligonucleotides for *GLC7* and snR44 were used (listed in Supporting Information 2).

**Real time PCR**: The expression levels of the *GLC7* gene in the intron replacement mutant and BY4742 wild type grown in F1 media and in F1 + 0.9M NaCl were assessed by quantitative real-time PCR using the QuantiTect real time PCR kit (Qiagen-catalogue no. 204143). cDNA was extracted using QuantiTect Reverse Transcription Kit (QIAGEN-catalogue no. 205311) according to manufacturer’s manual. Real time primers are listed in Supporting Information 2. The PCR reactions were performed in triplicate for two independent biological replicas, as described in previously (Naseeb and Delneri 2012). Relative normalized fold expression was calculated according to the ΔΔCt method using *ACT1* as a reference gene.

**RNAseq**: We used our previously-generated RNAseq data, deposited in GEO under the accession number GSE58884 (HOOKS *et al*. 2014). A total of 77286181 50 bp reads were filtered using the approach of Sassoon and Michael (2010). Filtering left 45520779 reads with an average quality > 20, which were mapped to the *S. cerevisiae* genome (SacCer3) using Bowtie with settings –m 1 –v 2 (Langmead *et al*. 2009; Trapnell *et al*. 2009). A total of 25254315 reads with a maximum of two mismatches were mapped in the *S. cerevisiae* genome. To calculate average number of reads per intron or CDS (in Reads Per Kilobase per Million mapped reads, RPKM), reads for each intron or CDS were summarized using *featureCounts* (Liao *et al*. 2014) and divided by the length of the feature in kB and the million reads mapped. The number of reads mapping to introns and their host genes is presented in Supporting Information 3.

**Analysis of exosome target data**: The CRAC data presented by Schneider *et al*. (2013) were filtered for genes that contain introns. From the RNAseq data presented here, we calculated the number of reads in RPKM corresponding only to the ORFs for the same set of genes. For each individual CRAC experiment, the number of reads for each gene was normalized by the number of ORF reads for this gene from our RNAseq data. The percentile rank of normalized values was calculated for each gene in each CRAC experiment, and then averaged across the sixteen CRAC experiments (Supporting Information 3). We considered genes with an average percentile rank with the top 10% to be preferentially bound by the exosome protein components.

**Deletion of predicted intronic RNA**: In order to generate deletions of intron fragments or insertions into introns, the Cre-loxP system was used with a kanMX cassette flanked by loxP sites (Guldener *et al*. 1996). Deletion cassettes were amplified as described previously (Delneri *et al*. 2003). The *S. cerevisiae* BY4743 strain was transformed with 1 µg of each PCR product according to Gietz and Schiestl (2007). Selection of mutants and PCR confirmation were performed as described by Carter and Delneri (2010). The strains with the loxP-kanMX-loxP cassette were then transformed with the plasmid containing Cre-recombinase to excise the sequence between two loxP sites. Cre-recombinase was induced by culturing the cells overnight in the YP-raffinose medium and then for 2-3 h in YP-galactose medium. KanMX excision was confirmed by PCR. Dissection of tetrads was performed using a Singer MSM 300 microdissector (Delneri *et al*. 2003). Haploids displaying the BY4742 metabolic background were chosen after series of cultures on SD solid medium lacking Lys, Met or Ura.

The effect of the loxP deletion was to replace the 204 bp (*GLC7* ncRNA deletion mutant) and 140 bp (*GLC7* control deletion mutant) intronic regions with 139 bp of the loxP scar with the fragments of the transformation vector. We also constructed an insertion mutant with the 139 bp remnant of the cassette inserted into the middle of the predicted RNA structure without deleting any intron bases. All primers used in creation of mutant strains are listed in Supporting Information 2.

**Growth rate assay**: Growth properties of the BY4742 strain and the intron mutants were assessed by time course growth profiles obtained using a FLUOstar optima microplate reader. Cells were cultured to stationary phase in YPD or F1 medium. The OD was measured at 595 nm and the cultures were diluted to an OD at 595 nm of 0.1 with pre-warmed YPD media, F1 media or F1 media containing 0.9 M NaCl. Each of the 96 plate wells was filled with 240 µl of diluted culture or media control. Absorbance measurements were taken every 5 minutes immediately after 1 minute shaking. Growth curves were plotted and analyzed using R according to a modified version of the method specified previously (Norris *et al*. 2013). In brief, the area under each growth curve (AUC) was calculated by the pracma package from normalized absorbance data. Additionally, we fitted growth curves from data points taken every 30 minutes using grofit R package with default settings (Kahm *et al*. 2010). Maximum growth rate, lag-phase and maximum growth were calculated from the fitted curves.

**Competitive growth test**: Competitive growth assays were performed in 8 ml media by adding equal number of cells (2 × 10^5^ cells/ml) of a mutant strain and the BY4742 reference strain, which had the *HO* gene replaced with kanMX as a marker to facilitate selection between strains. The *GLC7*/reference competition was performed in F1 media containing 0.9 M NaCl. The cultures were grown at 30 °C and maintained in log-phase by diluting each culture to 2 × 10^5^ cells/ml in fresh media every 12-24 h until generation number 37 ± 1 or 50 ± 2 was attained for F1 + 0.9 M NaCl or YPD media, respectively. The number of generations was calculated as described by Parenteau *et al*. (2008). When the appropriate generation number had been reached, approximately 200 cells were plated on YPD media and after 2 days replicated onto YPD + geneticin. Cells were counted to obtain the ratio of mutant versus reference strains.

## ACKNOWLEDGEMENTS

We are grateful to James Allen for assistance with EvoFold predictions. KBH was supported by a Wellcome Trust PhD studentship (grant number WT086809).

## AUTHOR CONTRIBUTIONS

KBH, SGJ and DD conceived and design the experiment. KBH performed the predictions, Northern, RT-PCRs, RNAseq and constructed the ncRNA *GLC7* mutant. SN performed the real-time PCR, constructed the *GLC7* negative control mutant and characterized the growth of all *GLC7* strains. KBH, SGJ and DD wrote the manuscript with input from SN.

## SUPPLEMENTARY INFORMATION

**Figure S1.**
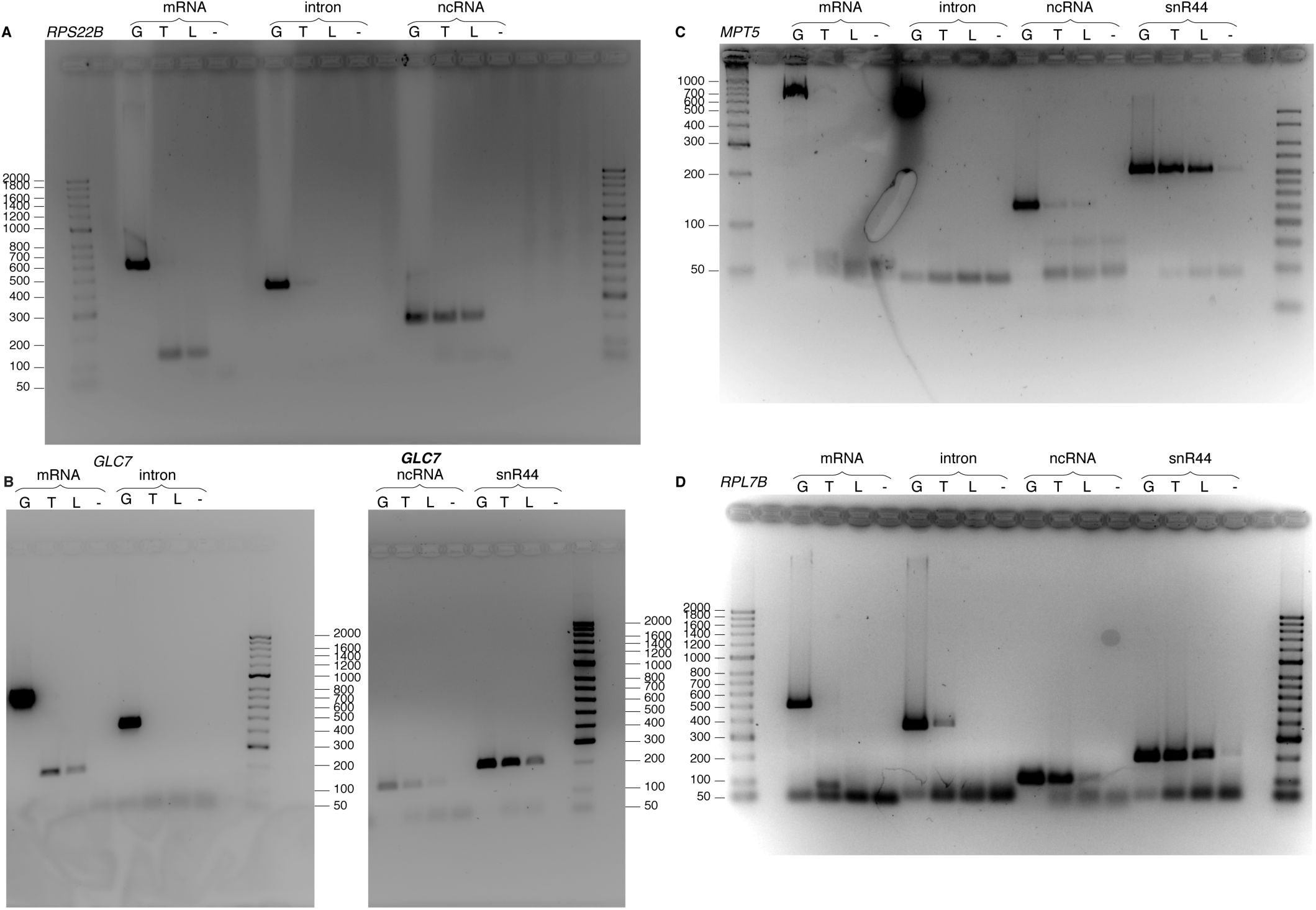
RT-PCR confirmation of ncRNA expressed from introns. Products for:

- A *RPS22B*
- B *GLC7*
- C *MPT5*
- D *RPL7B*. Lane designations for DNA templates: G, Genomic DNA; T, total cDNA; L, low molecular weight enriched cDNA; (-), no template negative control. Except *GLC7* ncRNA, which was run on a separate gel, for each gene brightness and contrast applied equally across the entire gel

**Figure S2.**
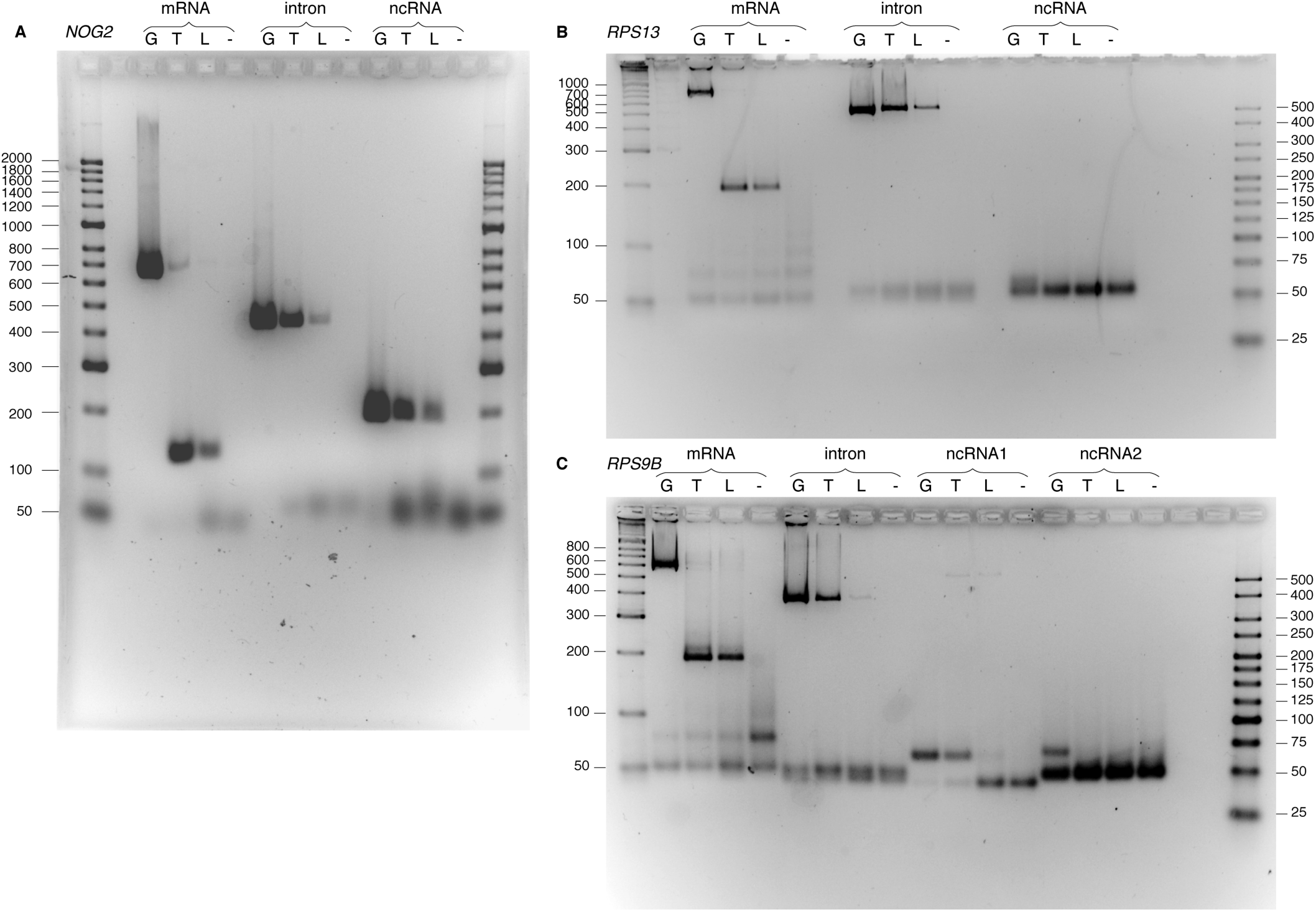
RT-PCR confirmation of intron expression accompanied by complete mRNA splicing. Products for:

- A *NOG2*
- B *RPS13*
- C *RPS9B*. Lane designations for DNA templates: G, genomic DNA; T, total cDNA; L, low molecular weight enriched cDNA; (-), no template negative control. Brightness and contrast were applied equally across each gel image.

**Figure S3.**
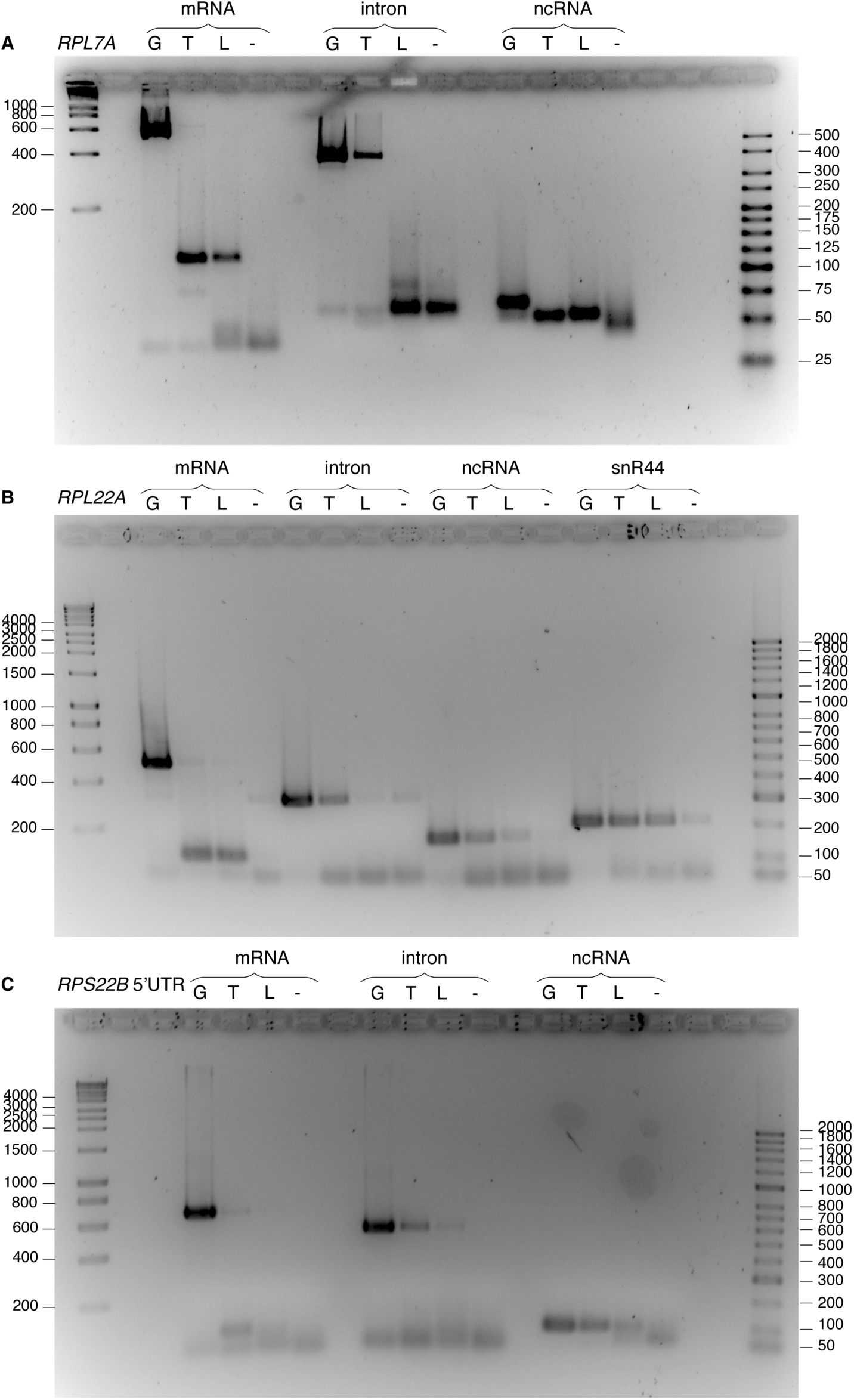
RT-PCR confirmation of intron expression accompanied by complete mRNA splicing. Products for: - A *RPL7A*
- B *RPL22A*
- C *RPS22B* 5‘UTR. Lane designations for DNA templates: G, genomic DNA; T, total cDNA; L, low molecular weight enriched cDNA; (-), no template negative control. Brightness and contrast were applied equally across each gel image.

**Figure S4.**
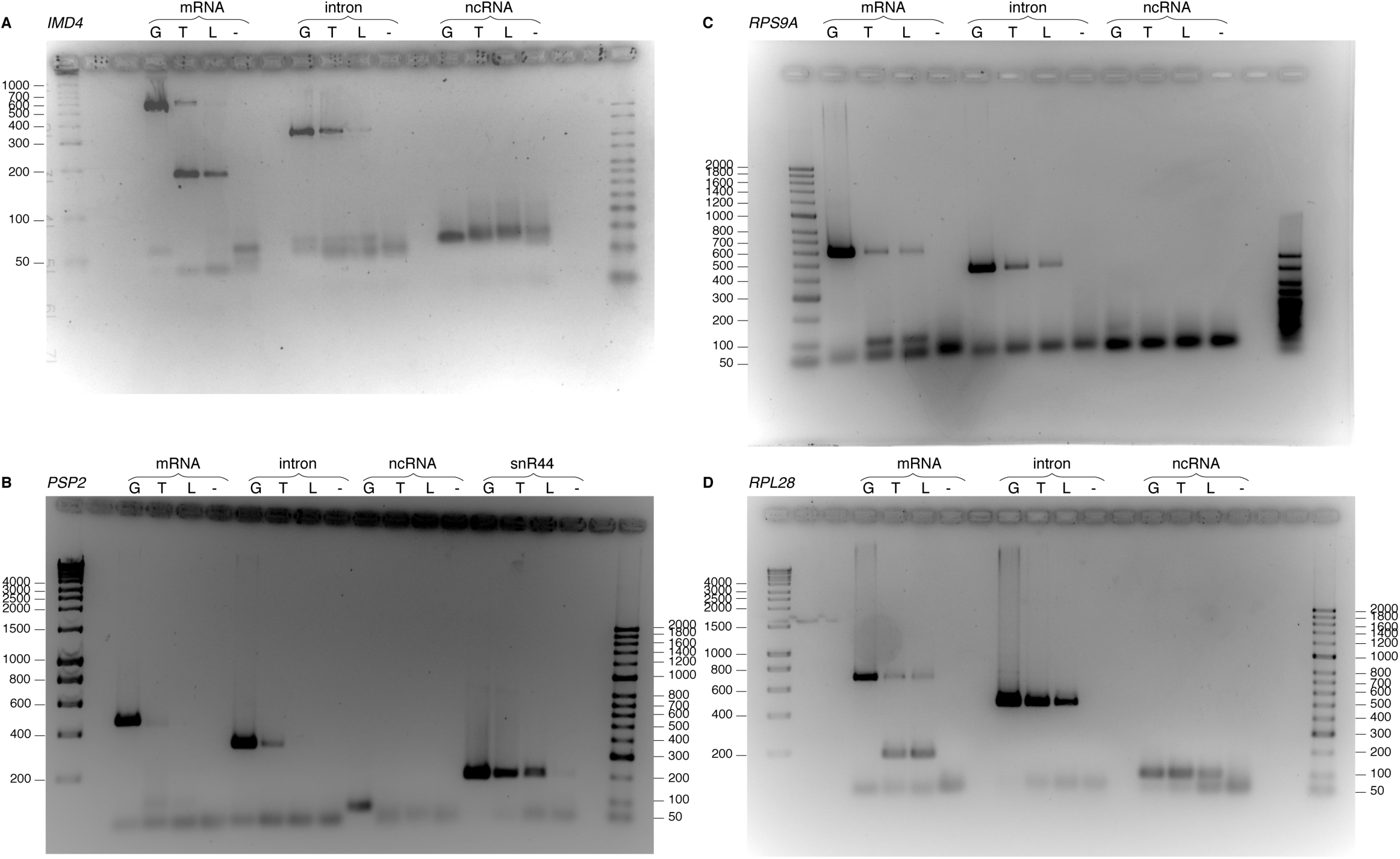
RT-PCR confirmation of alternative splicing. Products for:

- A *IMD4*
- B *PSP 2*
- C *RPS9A*
- D *RPL28*. Lane designations for DNA templates: G, genomic DNA; T, total cDNA; L, low molecular weight enriched cDNA; (-), no template negative

**Figure S5.**
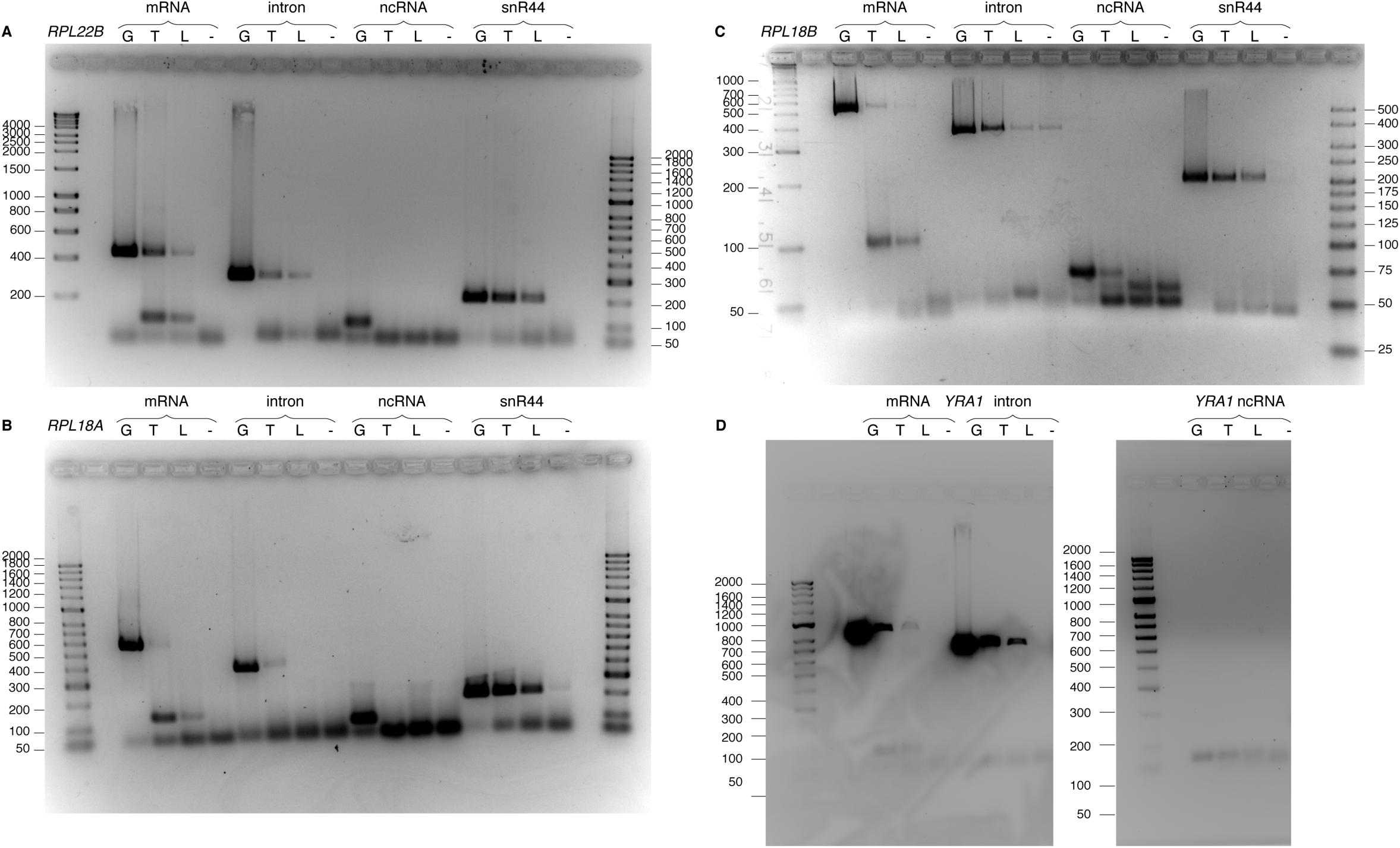
RT-PCR confirmation of alternative splicing. Products for:

- A *RPL22B*
- B *RPL18A*
- C *RPL18B*
- D *YRA1*. Lane designations for DNA templates: G, Genomic DNA; T, total cDNA; L, low molecular weight enriched cDNA; (-), no template

**Figure S6.**
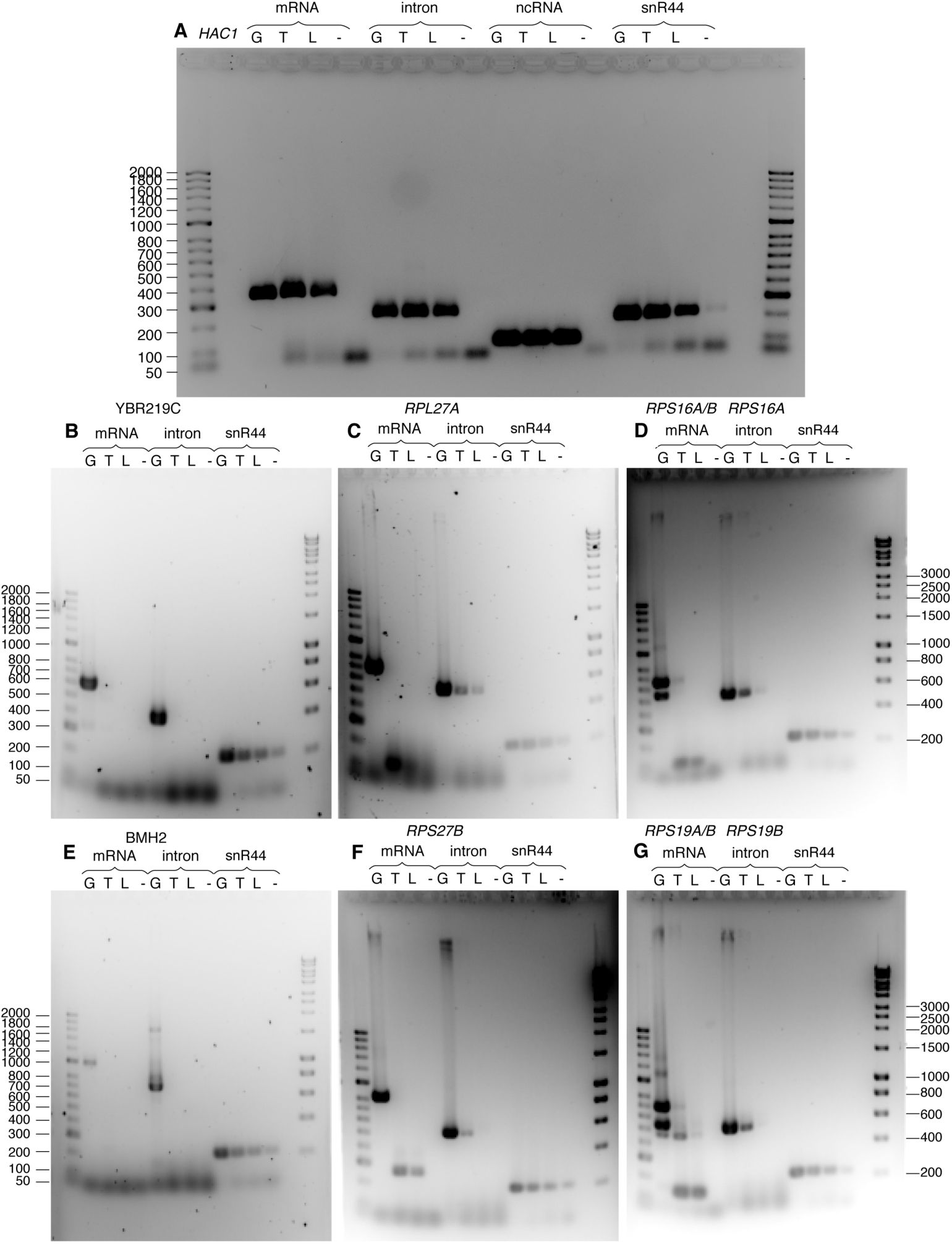
Control RT-PCRs confirming mRNA splicing status and intron expression. PCR products for genes: - A *HAC1*
- B YBR219C
- C *RPL27*
- D *RPS16A/B*
- E *BMH2*
- F *RPS27B*
- G *RPS19A/B*. Lane designations for DNA templates: G, Genomic DNA; T, total cDNA; L, low molecular weight enriched cDNA; (-), no template negative control. Brightness and contrast was applied equally across each gel image. For *RPS16A* and *RPS19B* designed mRNA primers also hybridize with the paralogous genes *RPS16B* and *RPS19A* giving two products from genomic DNA template.

**Figure S7.**
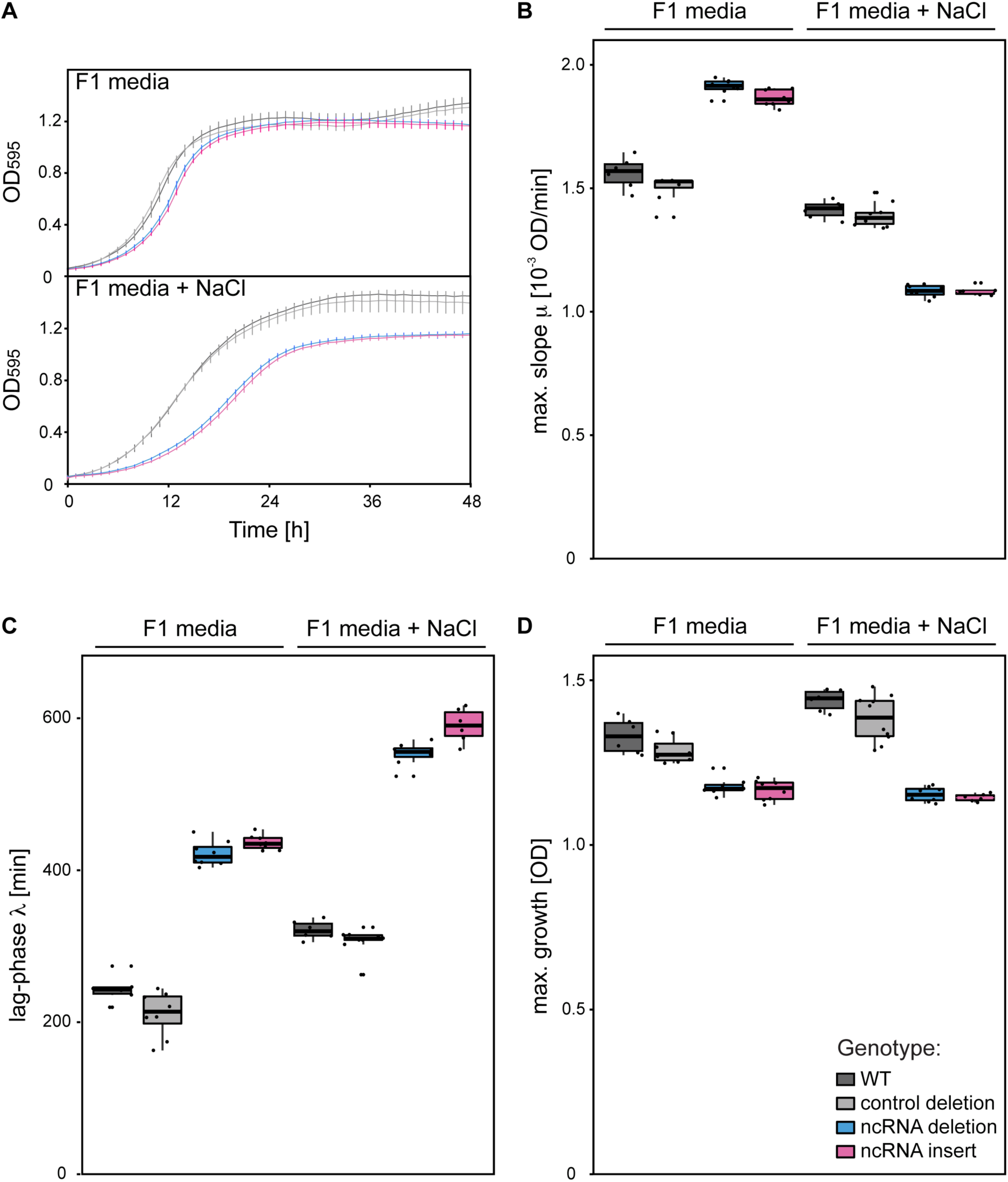
Growth of *GLC7* mutants in F1 and F1 + NaCl media. - A Growth curves made by plotting average OD measurements for each mutant taken every 60 minutes with bars representing standard deviation.
- B Maximum slope of a growth curve for each mutant.
- C Lag-phase time estimated from growth curves.
- D Maximal growth for each mutant. Mutants are labelled as displayed on the legend in the bottom right corner. Data presented in (B)-(D) was obtained by growth curve fitting with R package grofit.

**Table S1.** List of fungal genomes used for BLAST searches.

| Organism | Strain | Source | Ver | Reference<br>[PMID] |
| --- | --- | --- | --- | --- |
| *Aspergillus fumigatus* | Af293 | Broad Inst. | 1 | 16372009 |
| *Aspergillus nidulans* | FGSC A4 | Broad Inst. | 1 | 16372000 |
| *Aspergillus oryzae* | RIB40 | NITE | 1 | 16372010 |
| *Aspergillus terreus* | NIH2624 | Broad Inst. | 1 | - |
| *Botryotinia fuckeliana* | B05.10 | Broad Inst. | 1 | - |
| *Candida albicans* | SC5314 | Stanford Uni | 2 | 15123810 |
| *Candida glabrata* | CBS138 | Genolevures | 1 | 15229592 |
| *Chaetomium globosum* | CBS 148.51 | Broad Inst. | 1 | - |
| *Coccidioides immitis* | H538.4 | Broad Inst. | 1 | - |
| *Coprinopsis cinerea* | okayama7#130 | Broad Inst. | 1 | - |
| *Cryptococcus neoformans* | serA H99 | Broad Inst. | 1 | - |
| *Debaryomyces hansenii* | CBS767 | Genolevures | 1 | 15229592 |
| *Eremothecium gossypii* | ATCC 10895 | Biozentrum | 1 | 15001715 |
| *Fusarium graminearum* | PH-1 | Broad Inst. | 3 | 17823352 |
| *Fusarium verticillioides* | 7600 | Broad Inst. | 3 | - |
| *Histoplasma capsulatum* | G217B | WashU | 1 | - |
| *Kluyveromyces lactis* | NRRL Y-1140 | Genolevures | 1 | 15229592 |
| *Lachancea kluyveri* | NRRL Y-12651 | Stanford Uni | 1 | - |
| *Lachancea waltii* | NCYC 2644 | Broad Inst. | 1 | 15004568 |
| *Magnaporthe grisea* | 70-15 | Broad Inst. | 5 | 15846337 |
| *Naumovozyma castellii* | NRRL Y-12630 | Stanford Uni | 1 | 15229592 |
| *Neurospora crassa* | OR74A | Broad Inst. | 7 | 12712197 |
| *Parastagonospora nodorum* | SN15 | Broad Inst. | 1 | - |
| *Phanerochaete chrysosporium* | RP-78 | DOE Joint Genome | 1 | 15122302 |
| *Rhizopus oryzae* | RA 99-880 | Broad Inst. | 1 | - |
| *Saccharomyces bayanus* var. *uvarum* | 623-6C | Stanford Uni | 1 | 15229592 |
| *Saccharomyces cerevisiae* | S288c | SGD | 1 | 9169864 |
| *Saccharomyces kudriavzevii* | IFO 1802 | Stanford Uni | 1 | 15229592 |
| *Saccharomyces mikatae* | IFO 1815 | Stanford Uni | 1 | 15229592 |
| *Saccharomyces paradoxus* | NRRL Y-17217 | Stanford Uni | 1 | 15229592 |
| *Schizosaccharomyces japonicus* | yFS275 (SJ5) | Broad Inst. | 1 | 21511999 |
| *Schizosaccharomyces pombe* | 972h | Sanger Inst. | 1 | 11859360 |
| *Sclerotinia sclerotiorum* | 1980 | Broad Inst. | 1 | - |
| *Trichoderma reesei* | QM6a | DOE Joint | 2 | 18454138 |
| *Uncinocarpus reesii* | 1704 | Broad Inst. | 1 | - |
| *Ustilago maydis* | 521 | Broad Inst. | 1 | 17080091 |
| *Yarrowia lipolytica* | CLIB122 | Genolevures Cons. | 1 | 15229592 |

## SUPPLEMENTARY DATASETS

**Supplementary Dataset 1**. List of RNAz, CMfinder and EvoFold predictions for each intron.

**Supplementary Dataset 2**. Primer sequences and probes used in the study. List contains forward and reversed primers used for generating and confirming deletion mutants, RT-PCR and real time PCR.

**Supplementary Dataset 3**. Expression of introns and coding sequences of host genes [RPKM] and the average percentile from exosome targets data (CRAC) for each gene.

